## Supplementary Material for "Adult dentate gyrus neurogenesis: a functional model"

### Analytical computation of the L2-norm and angle

We consider the case where two mature DGCs have learned their synaptic connections, such that the first mature DGC with feedforward weight vector $\vec{w}_1$ is selective for cluster 1 with normalized center of mass $\vec{P}^1$ (see Methods), and the second mature DGC with feedforward weight vector $\vec{w}_2$ is selective for cluster 2 with normalized center of mass $\vec{P}^2$. After convergence, we have $\vec{w}_1 = \langle||\vec{w}_1||\rangle\vec{P}^1$ and $\vec{w}_2 = \langle||\vec{w}_2||\rangle\vec{P}^2$, where $\langle||\vec{w}_k||\rangle$ is the expected L2-norm of the feedforward weight vector onto mature DGC $k$ that is selective for pretrained cluster $k$. In addition, the upper bound for the L2-norm of the weight vectors of the mature DGCs can be determined $\langle||\vec{w}_1||\rangle = \langle||\vec{w}_2||\rangle \leqslant 1.5$ (Methods). In our case, we obtain $\langle||\vec{w}_1||\rangle = \langle||\vec{w}_2||\rangle \approx 1.49$ because of the dispersion of the patterns around their center of mass, hence we will use this value for the numerical computations below.

We represent the feedforward weight vector $\vec{w}_i$ onto a newborn DGC as an arrow of length $\langle||\vec{w}_1||\rangle$ (Suppl. Fig. S3e,f). We compute analytically its L2-norm at the end of the early phase of maturation of the newborn DGC, as well as its angle $\phi$ with the center of mass of the novel cluster $\vec{P}^i$, to confirm the results obtained numerically (Fig. 6 and Suppl. Fig. S3).

In the early phase of maturation, the feedforward weight vector onto the newborn DGC grows. The norm stabilizes at a higher value in the case of similar patterns ($s = 0.8$, Suppl. Fig. S3a) than in the case of distinct patterns ($s = 0.2$, Suppl. Fig. S3c). It is due to the fact that the center of mass of three *similar* clusters lies closer to the surface of the sphere than the center of mass of two *distinct* clusters (see below). In the late phase of maturation, for similar clusters we observe a slight increase of the L2-norm of the feedforward weight vector onto the newborn DGC concomitantly with the decrease of angle with the center of mass of the novel cluster (Suppl. Fig. S3b), because the center of mass of the novel cluster lies closer to the surface of the sphere than the center of mass of the three clusters.

#### Similar clusters

The angle between the center of mass of any pair of similar clusters ($s = 0.8$, $\xi = 0.2$) is given by equation (16):

$$\Omega_S = \arccos\left(\frac{1}{1+0.2^2}\right) \tag{31}$$

Half the distance between the projections of the center of mass of any pair of two similar clusters on a concentric sphere with radius $\langle ||\vec{w}_1|| \rangle$ is given by (Suppl. Fig. S3e):

$$z = \langle ||\vec{w}_1|| \rangle \cdot \sin\left(\frac{\Omega_S}{2}\right) \tag{32}$$

The triangle which connects the centers of masses of the three clusters is equilateral, and $y$ separates one of its angle in two equal parts ($\pi/6$ [rad] each). So the length $y$ can be calculated:

$$y = \frac{z}{\cos\left(\frac{\pi}{6}\right)} \tag{33}$$

Using Pythagoras formula, we can thus determine the expected L2-norm $\langle ||\vec{w}_i|| \rangle$ of the feedforward weight vector onto the newborn DGC at the end of the early phase of maturation:

$$\langle ||\vec{w}_i|| \rangle = \sqrt{\langle ||\vec{w}_1|| \rangle^2 - y^2}, \tag{34}$$

and finally its angle with the center of mass of the novel cluster:

$$\phi = \arccos\left(\frac{\langle ||\vec{w}_i|| \rangle}{\langle ||\vec{w}_1|| \rangle}\right) \tag{35}$$

The numerical values are: $\langle ||\vec{w}_i|| \rangle \approx 1.47$ and $\phi \approx 9.21[^\circ]$, which correspond to the values on Suppl. Fig. S3a.

### Distinct clusters

In the case of distinct patterns ($s = 0.2$, $\xi = 0.8$), the angle between the center of mass of any pair of clusters is given by equation (16):

$$\Omega_D = \arccos\left(\frac{1}{1 + 0.8^2}\right) > \Omega_S \tag{36}$$

We can directly compute the expected L2-norm of the feedforward weight vector onto the newborn DGC at the end of the early phase of maturation (Suppl. Fig. S3c):

$$\langle ||\vec{w}_i|| \rangle = \langle ||\vec{w}_1|| \rangle \cdot \cos\left(\frac{\Omega_D}{2}\right) \tag{37}$$

We can then calculate the length $z$ between the projection of the center of mass of one of the two pretrained clusters on a concentric sphere with radius $\langle ||\vec{w}_1|| \rangle$ and the feedforward weight vector onto the newborn DGC:

$$z = \langle ||\vec{w}_1|| \rangle \cdot \sin\left(\frac{\Omega_D}{2}\right) \tag{38}$$

Analogous to the similar case, we observe that $y$ separates one angle of the equilateral triangle connecting the projections of the center of mass of the clusters on the sphere in two equal parts, consequently:

$$y = \frac{z}{\tan\left(\frac{\pi}{6}\right)} \tag{39}$$

Finally, the angle between the center of mass of the novel cluster and the feedforward weight vector onto the newborn DGC at the end of the early phase of maturation is:

$$\phi = \arccos\left(\frac{\langle ||\vec{w}_i||\rangle^2 + \langle ||\vec{w}_1||\rangle^2 - y^2}{2\langle ||\vec{w}_i||\rangle\langle ||\vec{w}_1||\rangle}\right) \tag{40}$$

We obtain the following approximate values: $\langle ||\vec{w}_i||\rangle \approx 1.34$ and $\phi \approx 47.2[^\circ]$, which correspond to the values on Suppl. Fig. S3c. The angle $\phi$ is smaller in the similar case than in the distinct case, hence the norm is larger in the similar case, as observed in Suppl. Fig. S3a,c.

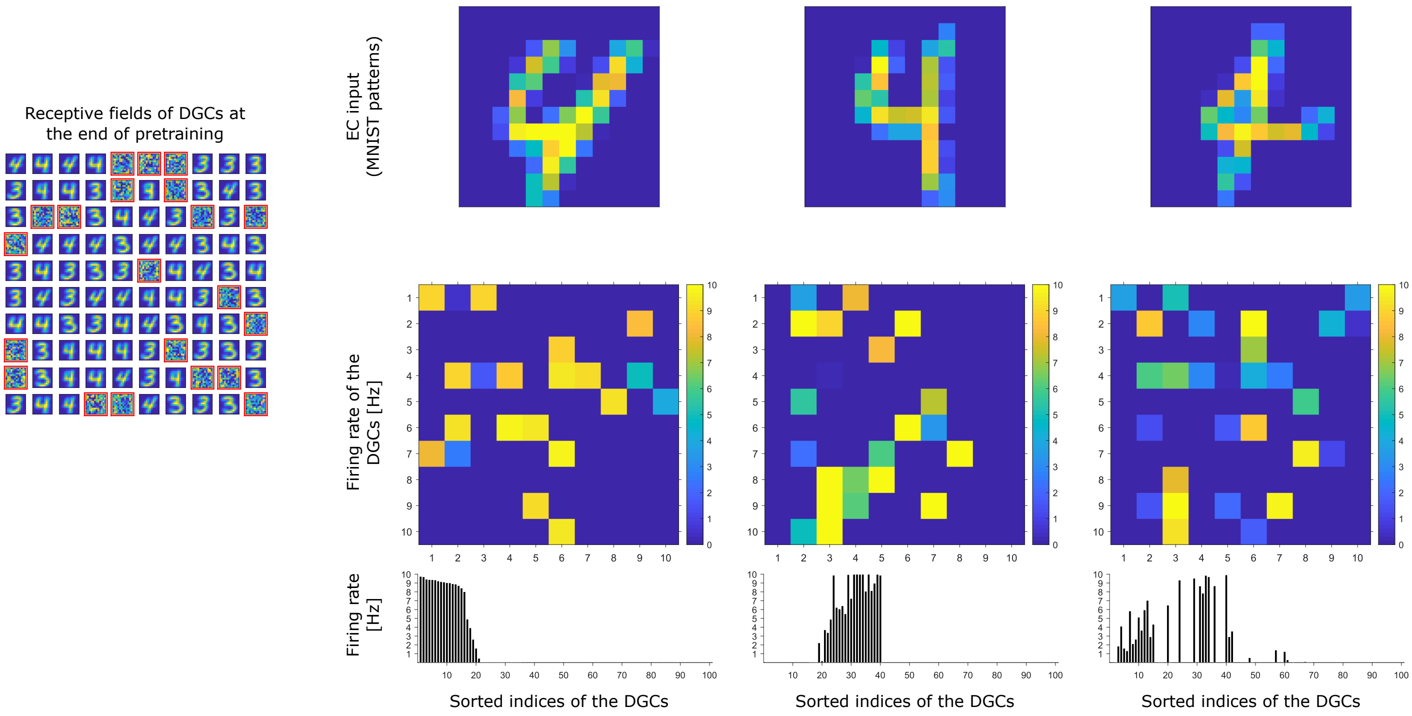


**Figure S1: Activity of 100 model DGCs in response to different patterns.** At the end of pretraining, three different patterns of digit 4 applied to EC neurons (top) cause different firing rate patterns of the 100 DGCs arranged in a matrix of 10-by-10 cells (middle). DGCs with a receptive field (left: 10-by-10 grid of receptive fields) similar to a presented EC activation pattern respond more strongly than the others. Bottom: Firing rates of the DGCs with indices sorted from highest to lowest firing rate in response to the first pattern. All 3 patterns shown come from the testing set, and are correctly classified using our readout network.

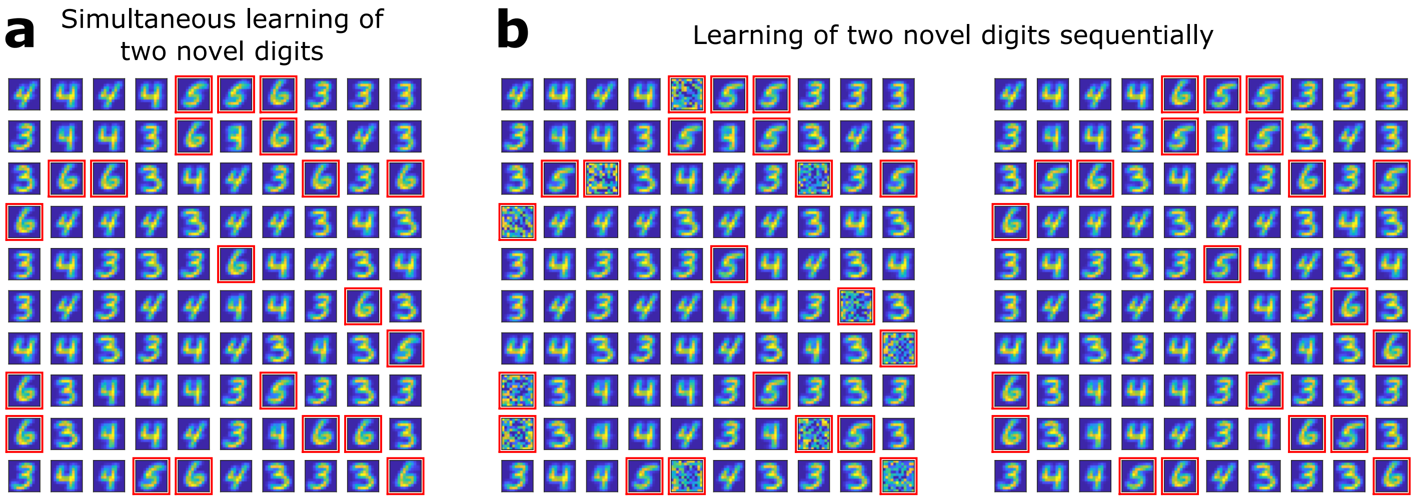


**Figure S2: Receptive fields of DGCs.** (**a**) Several novel digits can be learned simultaneously. After pretraining as in Fig. 1e, unresponsive neurons are replaced by newborn DGCs. When patterns from digits 3, 4, 5, and 6 are presented in random order, newborn DGCs exhibit after maturation receptive fields with selectivity for the novel digits 5 and 6. (**b**) Several novel digits can be learned sequentially. After pretraining with digits 3 and 4, ten randomly selected unresponsive neurons are replaced by newborn DGCs. Patterns from digits 3, 4, and 5 are presented in random order, while newborn DGCs mature and develop selectivity for the novel digit 5, with different writing styles. Later, the eleven remaining unresponsive neurons of the network are replaced by newborn DGCs. When patterns from the novel digit 6 are presented intermingled with patterns from digits 3, 4, and 5, the newborn DGCs develop selectivity for digit 6.

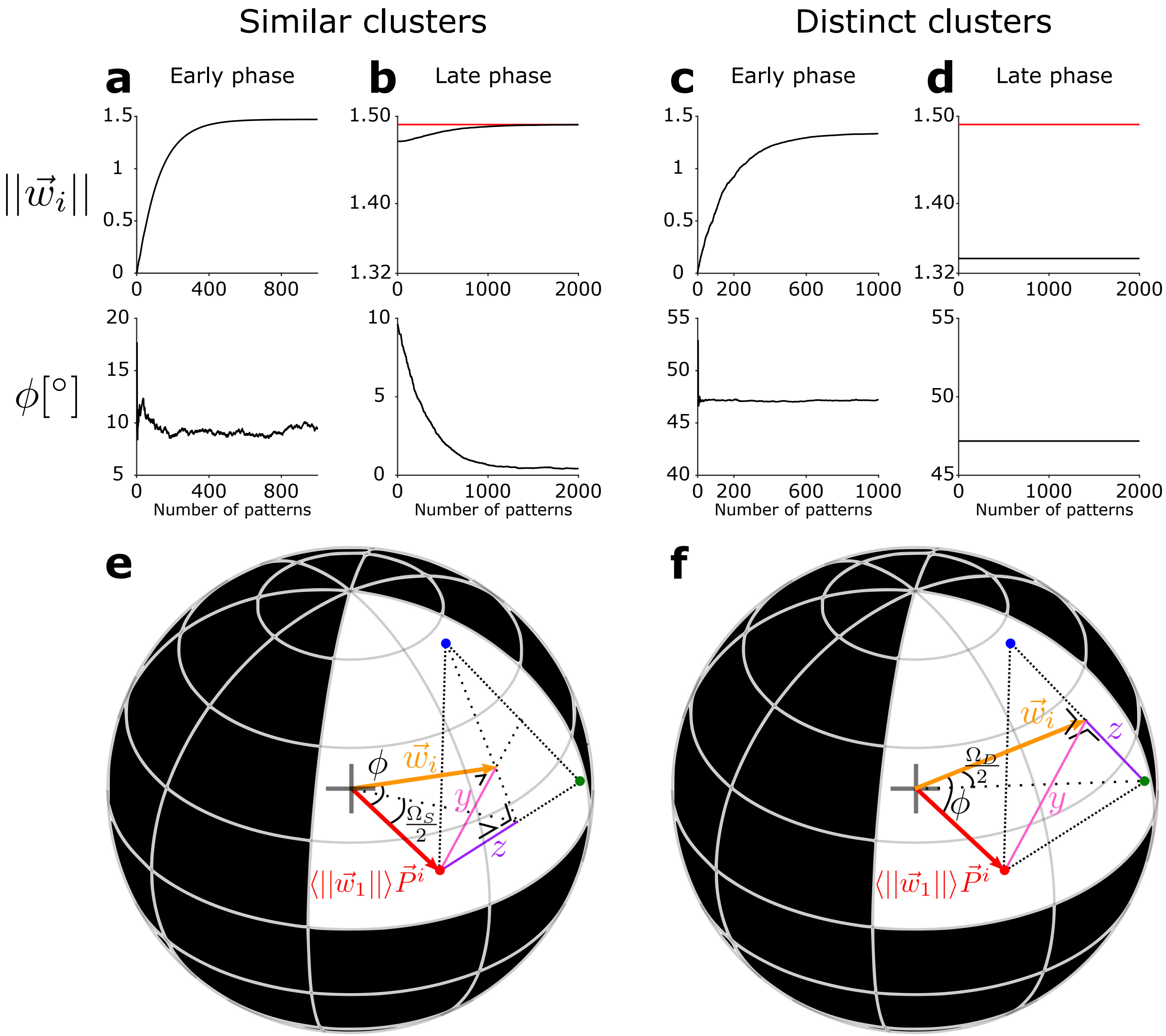


**Figure S3: Evolution of the feedforward weight vector onto the newborn DGC.** (**a-d**) The total synaptic strength $||\vec{w}_i|| = \sqrt{\sum_j (w_{ij})^2}$ of the weight vector onto the newborn DGC (top row), and of its angular separation $\phi$ with the center of mass of the novel cluster (bottom row), as a function of the number of pattern presentations. (**a**,**b**) The three clusters are similar ($s = 0.8$). (**c**,**d**) The three clusters are distinct ($s = 0.2$). Early phase of maturation (**a**,**c**), late phase of maturation (**b**,**d**). The red line shows the mean value of the synaptic strength of the mature DGCs. (**e**,**f**) Schematic drawing for the analytical computations. The L2-norm of the weight vector $\vec{w}_i$ onto the newborn DGC at the end of the early phase of maturation, and its angle $\phi$ with the center of mass of the novel cluster, for (**e**) similar clusters ($s = 0.8$), and (**f**) distinct clusters ($s = 0.2$). The sphere has a radius $\langle||\vec{w}_1||\rangle$. The centers of mass of the first two clusters (represented by the two mature DGCs) projected onto the hypersphere are represented by the blue and green dots. The red dot represents the projection of the center of mass of the novel cluster, $\vec{P}^i$, onto the hypersphere.

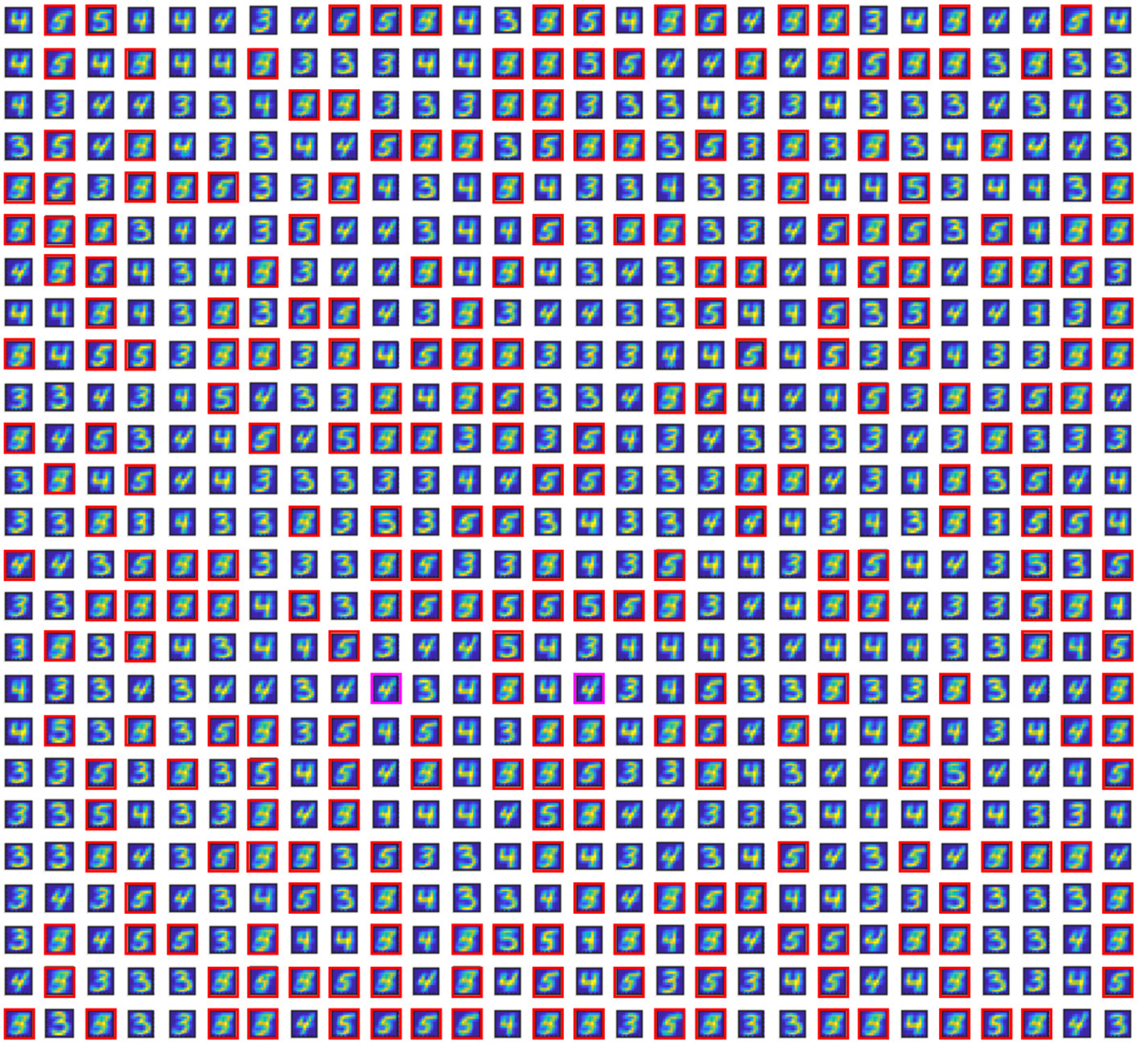

**Figure S4: Receptive fields of the DGCs in a larger network with $N_{DGC} = 700$ (all other parameters unchanged).** After pretraining with digits 3 and 4, all 275 unresponsive DGCs (highlighted by the red/magenta squares) are replaced by newborn DGCs. Newborn DGCs follow a two-step maturation process while patterns from digits 3, 4 and 5 are presented to the network. At the end of maturation, most newborn DGCs represent different prototypes of the novel digit 5. Two of them (highlighted in magenta) became selective for digit 4.

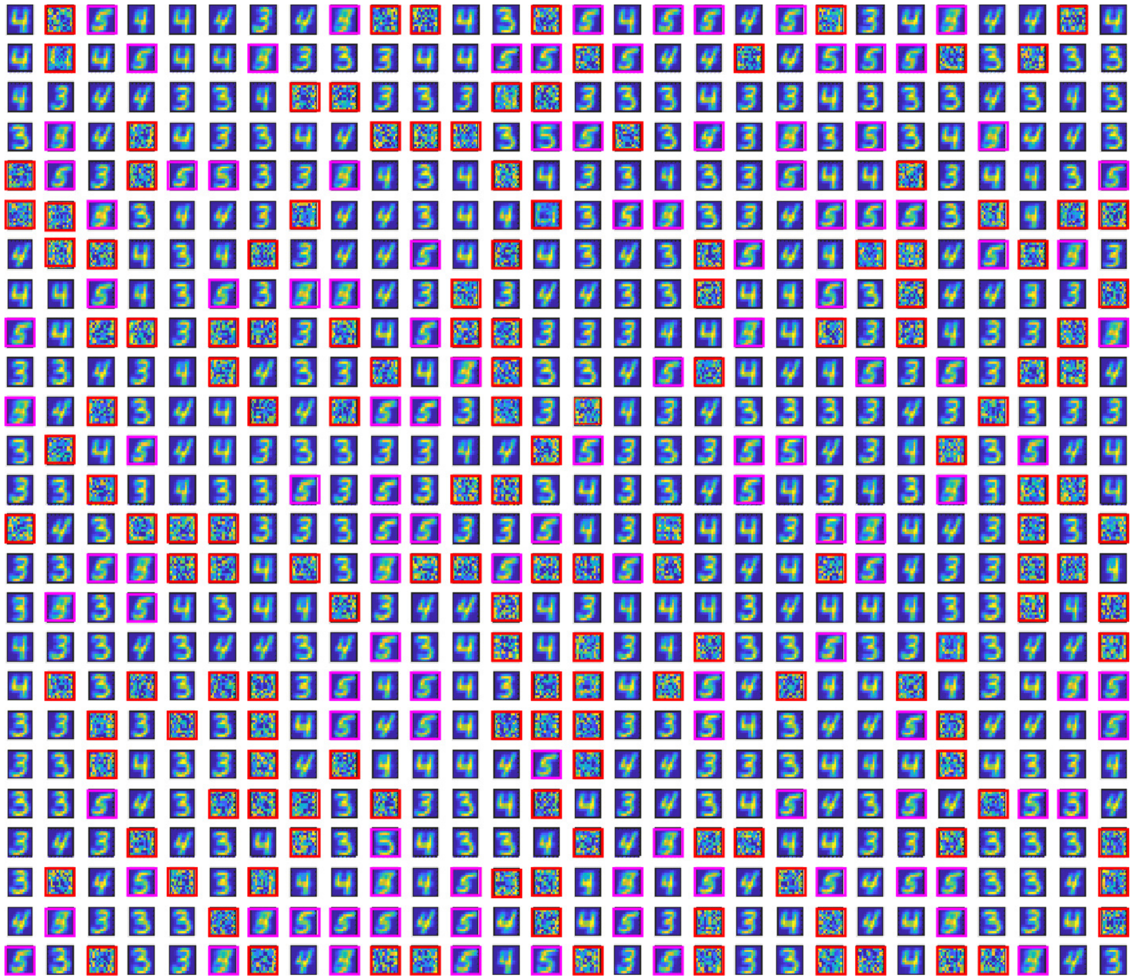

**Figure S5: Receptive fields of the DGCs in a larger network with $N_{DGC} = 700$ (all other parameters unchanged), when only a fraction of unresponsive units are replaced by newborn DGCs.** Out of the 275 unresponsive DGCs after pretraining with digits 3 and 4 (highlighted by the red/magenta squares), 119 are replaced by newborn DGCs (highlighted by the magenta squares), to mimic the experimental observation that only a fraction of DGCs are newborn DGCs. Newborn DGCs follow a two-step maturation process while patterns from digits 3, 4 and 5 are presented to the network. At the end of maturation, newborn DGCs represent different prototypes or features of the novel digit 5. The remaining unresponsive units (highlighted by the red squares) are available to be replaced later by newborn DGCs so as to learn further tasks.

**Table S1: Classification performance for random combinations of digits.** The classification performance ($P_0$, $P_1$, $P_2$) is defined as the percentage of correctly classified patterns on the test set. The numbers m n + q (first column) indicate that MNIST digits m and n are used for pretraining (second column); m, n and q are used for pretraining (third column); or m and n are used for pretraining, and patterns from digit q added after neurogenesis (fourth column). $P_2 - P_1$ (last column) is used for evaluating the contribution of neurogenesis to classification performance.

| Digits | Pretraining with two digits | Pretraining with three digits | Neurogenesis after pretraining with two digits | $P_2 - P_1$ |
|---|---|---|---|---|
| 3 4 + 5 | $P_0$ = 99.25% (digit 3: 98.71%; digit 4: 99.80%) | $P_1$ = 92.09% (digit 3: 86.83%; digit 4: 98.78%; digit 5: 90.70%) | $P_2$ = 94.56% (digit 3: 90.5%; digit 4: 98.17%; digit 5: 95.18%) | 2.47% |
| 6 8 + 2 | $P_0$ = 98.40% (digit 6: 98.02%; digit 8: 98.77%) | $P_1$ = 95.04% (digit 2: 94.77%; digit 6: 96.45%; digit 8: 93.94%) | $P_2$ = 96.12% (digit 2: 94.28%; digit 6: 97.81%; digit 8: 96.41%) | 1.08% |
| 0 1 + 6 | $P_0$ = 99.86% (digit 0: 99.80%; digit 1: 99.91%) | $P_1$ = 97.04% (digit 0: 94.90%; digit 1: 99.74%; digit 6: 96.03%) | $P_2$ = 97.20% (digit 0: 97.76%; digit 1: 99.38%; digit 6: 94.05%) | 0.16% |
| 2 9 + 4 | $P_0$ = 98.53% (digit 2: 98.84%; digit 9: 98.22%) | $P_1$ = 77.11% (digit 2: 97.58%; digit 4: 42.67%; digit 9: 89.69%) | $P_2$ = 89.84% (digit 2: 97.19%; digit 4: 78.72%; digit 9: 93.16%) | 12.73% |
| 4 7 + 6 | $P_0$ = 98.56% (digit 4: 99.19%; digit 7: 97.96%) | $P_1$ = 96.63% (digit 4: 94.40%; digit 6: 97.91%; digit 7: 97.57%) | $P_2$ = 97.61% (digit 4: 96.84%; digit 6: 98.33%; digit 7: 97.67%) | 0.98% |
| 5 9 + 3 | $P_0$ = 97.26% (digit 5: 97.53%; digit 9: 97.03%) | $P_1$ = 85.19% (digit 3: 84.55%; digit 5: 72.98%; digit 9: 96.63%) | $P_2$ = 93.47% (digit 3: 89.11%; digit 5: 95.07%; digit 9: 96.43%) | 8.28% |
| 4 9 + 6 | $P_0$ = 68.91% (digit 4: 53.05%; digit 9: 84.34%) | $P_1$ = 75.35% (digit 4: 35.44%; digit 6: 98.43%; digit 9: 92.27%) | $P_2$ = 77.45% (digit 4: 49.39%; digit 6: 98.23%; digit 9: 85.03%) | 2.10% |
| 6 7 + 3 | $P_0$ = 99.55% (digit 6: 99.37%; digit 7: 99.71%) | $P_1$ = 97.23% (digit 3: 95.35%; digit 6: 99.06%; digit 7: 97.37%) | $P_2$ = 97.66% (digit 3: 95.74%; digit 6: 99.27%; digit 7: 98.05%) | 0.43% |
| 0 3 + 2 | $P_0$ = 99.50% (digit 0: 99.59%; digit 3: 99.41%) | $P_1$ = 95.93% (digit 0: 98.27%; digit 2: 93.60%; digit 3: 96.04%) | $P_2$ = 96.56% (digit 0: 99.39%; digit 2: 93.90%; digit 3: 96.53%) | 0.63% |
| 3 5 + 8 | $P_0$ = 91.43% (digit 3: 86.83%; digit 5: 96.64%) | $P_1$ = 78.03% (digit 3: 81.58%; digit 5: 87.33%; digit 8: 65.81% | $P_2$ = 88.80% (digit 3: 87.43%; digit 5: 93.72%; digit 8: 85.73%) | 10.77% |

**Table S2: Comparison of networks with different numbers of inhibitory neurons.** The number of excitatory neurons is $N_{DGC} = 100$ for all three networks, and there are $N_I$ inhibitory neurons. The case with $N_I = 25$ is the one presented in the main text. All other network parameters are unchanged (including $p^*$). Each network is pretrained once with digits 3 and 4. The percentage of active neurons (firing rate $>$ 1 Hz) for each testing pattern of the corresponding digit is given (mean $\pm$ standard deviation), as well as the classification performance over all testing patterns from the trained digits.

| | | $N_I = 12$ | $N_I = 25$ | $N_I = 50$ |
|---|---|---|---|---|
| Percentage of unresponsive neurons at the end of pretraining | | 22% | 21% | 22% |
| Number of active neurons at the end of pretraining ($\mu \pm \sigma$) | Digit 3 | $22.05 \pm 4.18$ | $22.70 \pm 4.56$ | $22.42 \pm 4.54$ |
| | Digit 4 | $21.66 \pm 4.58$ | $22.09 \pm 3.91$ | $21.65 \pm 4.13$ |
| Number of active neurons at the end of the late phase of maturation ($\mu \pm \sigma$) | Digit 3 | $23.81 \pm 4.50$ | $24.16 \pm 4.87$ | $24.37 \pm 4.98$ |
| | Digit 4 | $22.38 \pm 4.92$ | $22.84 \pm 4.21$ | $22.41 \pm 4.46$ |
| | Digit 5 | $22.18 \pm 4.07$ | $21.98 \pm 4.52$ | $22.02 \pm 4.55$ |
| Classification performance at the end of pretraining | | 99.10% | 99.25% | 99.05% |
| Classification performance at the end of the late phase of maturation | | 94.17% | 94.56% | 94.45% |

**Table S3: Network with 700 DGCs (expansion factor from EC to dentate gyrus of about 5) compared to the case with $N_{DGC} = 100$ as in the main text.** All other network parameters are unchanged. Each network is pretrained with digits 3 and 4. Note that only a subset of neurons responsive to digit 3 (or 4) get active (firing rate $> 1$ Hz) for a given pattern 3 (or 4). Classification performance is evaluated over all test patterns from the trained digits. Top: after pretraining; bottom: late phase, after adding patterns from digit '5'. Either all unresponsive cells (Suppl. Fig. S4), or only a fraction of these (Suppl. Fig. S5), have been replaced by newborn model cells. For the network with 700 DGCs, about $16-18\%$ of DGCs are activated upon presentation of a digit 3 or 4 or 5 (about $112-126$ model DGCs). If 119 newborn DGCs are plastic during presentation of the novel digit 5 (middle column), these can become selective for prototypes of digit 5 (Suppl. Fig. S5) yielding a good classification performance while keeping 156 unresponsive DGCs available for future tasks. If only 35 newborn DGCs are available, classification performance is lower (right column).

| | | $N_{DGC} = 100$ | $N_{DGC} = 700$ | | |
|---|---|---|---|---|---|
| Percentage [in %] (and absolute number) of unresponsive neurons at the end of pretraining | | 21 (21 DGCs) | ~39 (275 DGCs) | | |
| Percentage [in %] (and absolute number) of selective neurons at the end of pretraining | Digit 3<br>Digit 4 | 40 (40 DGCs)<br>39 (39 DGCs) | ~30 (212 DGCs)<br>~30 (213 DGCs) | | |
| Percentage of active neurons per test pattern at the end of pretraining ($\mu \pm \sigma$ [%]) | Digit 3<br>Digit 4 | $22.70 \pm 4.56$<br>$22.09 \pm 3.91$ | $16.55 \pm 2.49$<br>$16.23 \pm 2.42$ | | |
| Classification performance at the end of pretraining [%] | | 99.25 | 99.55 | | |
| Percentage [in %] (and absolute number) of newborn model DGCs | | 21 (21 DGCs) | ~39 (275 DGCs) | 17 (119 DGCs) | 5 (35 DGCs) |
| Percentage of active neurons per test pattern at the end of the late phase of maturation ($\mu \pm \sigma$ [%]) | Digit 3<br>Digit 4<br>Digit 5 | $24.16 \pm 4.87$<br>$22.84 \pm 4.21$<br>$21.98 \pm 4.52$ | $17.56 \pm 2.73$<br>$16.56 \pm 2.57$<br>$16.62 \pm 2.85$ | $17.09 \pm 2.65$<br>$16.44 \pm 2.54$<br>$16.96 \pm 2.65$ | $16.67 \pm 2.53$<br>$16.30 \pm 2.45$<br>$17.78 \pm 2.56$ |
| Classification performance at the end of the late phase of maturation [%] | | 94.56 | 96.32 | 95.49 | 90.88 |
